# Functional characterization and optimization of protein expression in *Treponema denticola* shuttle plasmids

**DOI:** 10.1101/2024.10.27.620309

**Authors:** M. Paula Goetting-Minesky, Valentina Godovikova, Prakaimuk Saraithong, Alexander H. Rickard, Brigette R. Crawley, Sara M. Agolli, J. Christopher Fenno

**Affiliations:** Department of Biologic and Materials Sciences and Prosthodontics, School of Dentistry, University of Michigan, Ann Arbor, MI 48109; Department of Epidemiology, School of Public Health, University of Michigan, Ann Arbor, MI 48109

## Abstract

Oral spirochetes are among a small group of keystone pathogens contributing to dysregulation of periodontal tissue homeostasis, leading to breakdown of the tissue and bone supporting the teeth in periodontal disease. Of the greater than sixty oral *Treponema* species and phylotypes, *T. denticola* is one of the few that can be grown in culture and the only one in which genetic manipulation has been shown to be practicable. *T. denticola* is thus a model organism for studying spirochete metabolic processes, interactions with other microbes and host cell and tissue responses relevant to oral diseases as well as venereal and nonvenereal treponematoses. We recently demonstrated enhanced transformation efficiency using a SyngenicDNA-based shuttle plasmid resistant to *T. denticola* restriction-modification systems. Here we expand on this work by further characterizing the shuttle plasmid and optimizing expression of cloned genes using several promoter-gene constructs for genetic complementation and exogenous gene expression, including the first inducible system for controlled expression of potentially toxic plasmid-encoded genes in *Treponema*. Our results highlight the importance of precise pairing of promoters and genes of interest to obtaining biologically optimal protein expression. This work expands the utility of the shuttle plasmid and will facilitate future studies employing shuttle plasmids in analysis of *Treponema* physiology and behavior.

**IMPORTANCE:** Rigorous genetic analysis in oral spirochetes has been hampered by the limited utility of available versions of the *E. coli-T. denticola* shuttle plasmid system. We report expanded characterization of the shuttle plasmid, including relative activity of diverse promoters and the first inducible expression system described for *T. denticola.* We show that careful customization of the shuttle plasmid for specific applications is crucial for obtaining successful results.

## INTRODUCTION

*Treponema denticola*, the most readily cultivable of more than sixty oral *Treponema* species and phylotypes, is recognized as one of a few keystone pathogens driving periodontal disease development (1–4). While oral spirochetes are members of the commensal oral microbiome, their numbers are typically very low (or below detection levels) in health. They proliferate as periodontal disease develops, often comprising greater than 10% of total bacteria in diseased subgingival plaque (5). Both mechanical and antimicrobial treatment of periodontal disease result in clinical improvement and large reductions in spirochete levels (6, 7). As with other putative periodontal pathogens and pathobionts, the specific molecular mechanisms by which spirochetes contribute to disease are the subject of intensive studies by several groups.

Oral *Treponema* present unique challenges and opportunities to the field of molecular and cellular microbiology. These nutritionally fastidious anaerobes share with other spirochetes a cellular structure and motility apparatus unique among prokaryotes (8, 9). As both commensal residents and pathogens of the subgingival mucosa, *T. denticola* offers a wide range of potentially useful targets for research into microbe-host interactions and signaling, microbial communities, microbial physiology and molecular evolution. While pathogenic behaviors and disease associations of oral spirochetes are clearly distinct from those of *T. pallidum*, *T. denticola* is of significant interest as a model for the study of *Treponema* physiology, biosynthetic pathways and microbe-host interactions relevant to both human and animal treponemal diseases (3, 10, 11). This is due both to physiological similarities among *Treponema spp.* and to the fact that genetic studies in *T. pallidum* remain barely practicable due to limitations of its nascent systems for *in vitro* culture (12) and genetic manipulation (13).

Studies of the role of oral spirochetes in periodontal disease have been hampered both by difficulties in culturing these nutritionally fastidious anaerobes and by the limitations of available genetic technologies (14, 15). Allelic replacement mutagenesis of *T. denticola* was first reported over 20 years ago in ATCC 35405, the most highly studied strain (16). An *E. coli-T. denticola* shuttle plasmid was reported that functioned in *T. denticola* ATCC 33520 (17), but it could not be introduced into the most highly studied strain, ATCC 35405 (18, 19). *T. denticola* shuttle plasmids are all derived from cryptic plasmid pTS1 found in an oral *Treponema* clinical isolate (20) that was fused to an *E. coli* plasmid replicon to form shuttle plasmid (17). Plasmids based on this design have been used to complement mutations in *T. denticola* ATCC 33520 flagellar genes (17, 19, 21) and to express a *T. pallidum* lipoprotein in *T. phagedenis* (22), but plasmid-mediated mutant complementation was regarded as unfeasible in ATCC 35405, the most widely studied *T. denticola* strain. This has contributed to a lack of rigor in many *T. denticola* mutagenesis studies that did not include genetic complementation of *T. denticola* mutant strains.

There have been several notable recent advances in oral *Treponema* genetics including expansion of the number of available selectable markers in *T. denticola* (21, 23–26) and proofs of concept for transposon mutagenesis (27, 28). We previously reported improvements in genetic transformation methodology for *T. denticola* that included modification of shuttle plasmid pBFC (21) to remove a previously documented recognition site for Type II restriction enzyme TdeIII (18). This, combined with attention to appropriate methylation status of transforming plasmid DNA, facilitated routine transformation of *T. denticola* strains ATCC 35405 and ATCC 33520 with the modified shuttle plasmid pCF693 (29) (Table 1). More recently we characterized *T. denticola* restriction-modification (R-M) systems in two widely studied strains and demonstrated greatly enhanced transformation efficiency using a SyngenicDNA shuttle plasmid resistant to *T. denticola* R-M systems (28).

**TABLE 1.**
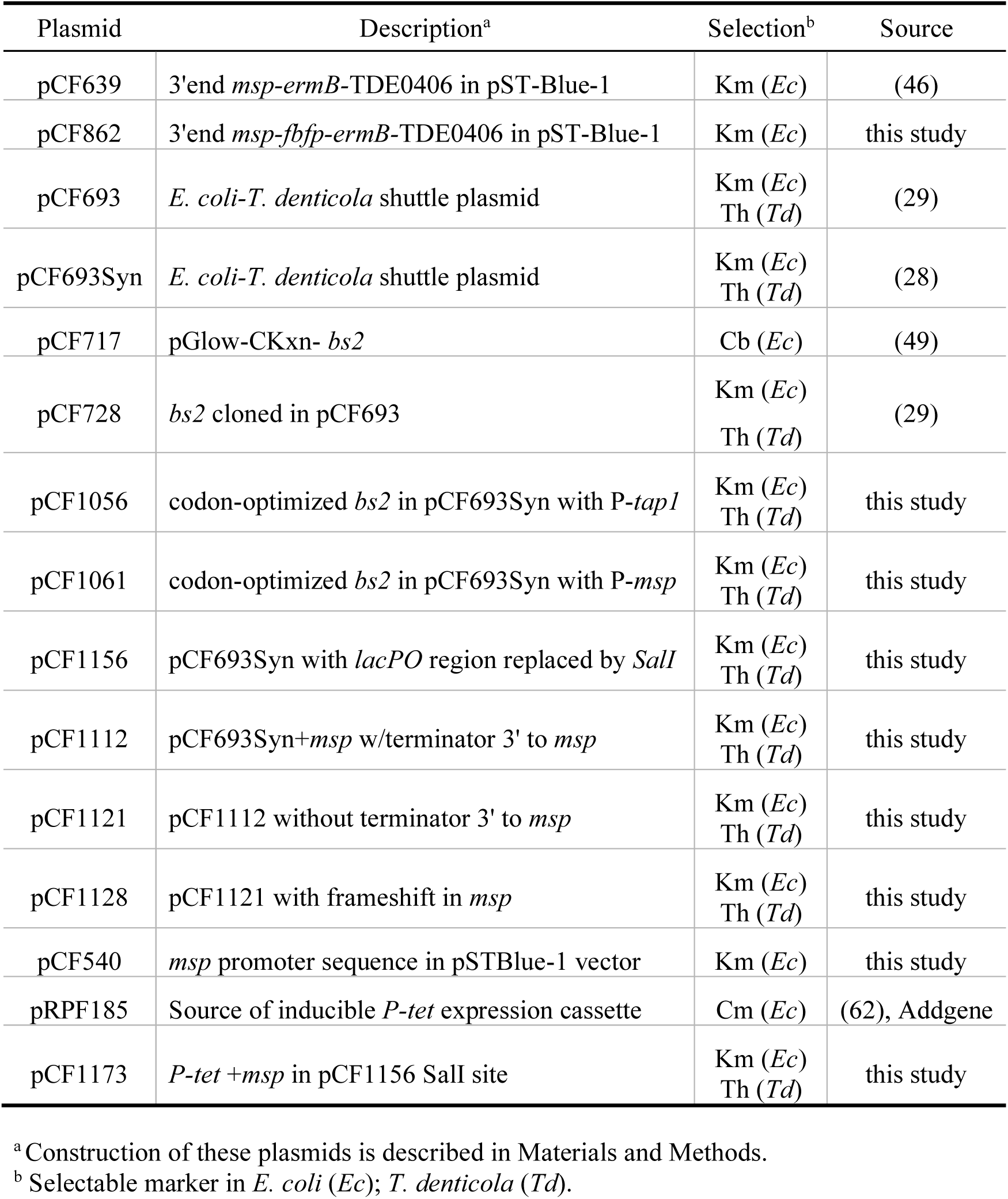
Plasmids used in this study.

Key factors that must be considered when designing plasmids for complementation experiments or for expression of heterologous genes include plasmid copy number, promoter activity and the nature of the cloned protein being expressed. Herein we report further characterization of the *T. denticola* shuttle plasmid system, including effects of codon optimization and promoter choice on expression of genes cloned in the shuttle plasmid. Finally, we report use of a tunable, tetracycline-inducible (iTet) promoter in the shuttle plasmid for genetic complementation of an isogenic mutant defective in the *T. denticola* major surface protein (Msp). Complementation of the *T. denticola* Λ*msp* strain demonstrates the utility of this tunable shuttle plasmid system in analysis of *T. denticola* outer membrane proteins whose overexpression is problematic in both *E. coli* and *T. denticola*.

## RESULTS AND DISCUSSION

### *T. denticola* native plasmids and shuttle plasmids

Two distinctly different native plasmids have been identified in oral *Treponema spp. T. denticola* ATCC 33520 carries a cryptic 2.65 kb plasmid pTD1 ((30); GenBank M87856) that is not found in *T. denticola* ATCC 35405. Plasmid pTD1 appears to replicate by a “rolling-circle” mechanism identified in bacteriophage and some Gram-positive bacterial plasmids (31, 32). The other plasmid (pTS1) was originally reported in an oral *Treponema* clinical isolate (20) and is the basis for the *E. coli-T. denticola* shuttle plasmid pKMR4PE (17), from which all subsequent *T. denticola* shuttle plasmids are derived. DNA sequences with high homology to pTS1 from at least three species of oral *Treponema* are present in Genbank. Other than pTS1 (Genbank AF112856), a complete plasmid sequence (pTPu1; Genbank CP009229) has been reported only in *T. putidum* OMZ 758 and it is identical to pTS1 (33).

Using oligonucleotide primers directed to the replication genes of pTD1 or pTS1 (Table S1), we screened 72 *T. denticola* strains (including more than 60 clinical isolates (34)) for the presence of either plasmid. None of the strains tested were positive in PCR for the pTS1 *rep* gene, and only two strains other than the *T. denticola* ATCC 33520 control were positive for the pTD1 *rep* gene (data not shown). This indicates that native plasmids are quite rare in *T. denticola.* We compared the sequence of pTD1 deposited in Genbank (M87856), with DNA sequences obtained by whole plasmid sequencing of pTD1 directly isolated from ATCC 33520 and the pTD1 homolog from a clinical isolate. Both plasmids contained a three-base insertion of an arginine residue (CGC) in the pTD1 *rep* gene compared with the original Genbank pTD1 sequence. Otherwise, we found only a very few minor differences (single base changes, no stop codon insertions) between all three sequences (Supplemental Material). These single base changes are possibly attributable to the inherent error rate observed in whole plasmid sequencing (<2%; Oxford Nanopore).

As a first step toward a rational strategy for optimized protein expression from the shuttle plasmid, we used quantitative PCR (qPCR) to determine the copy number of pCF728, the pCF693-derived shuttle plasmid carried in *T. denticola* CF734 (ATCC 35405/pCF728; Tables 1, 2) (29). Comparing the single-copy chromosomal gene *flaA* (TDE1712) with the gene encoding the Bs2 fluorescent protein (35, 36) carried on pCF728, we estimated the plasmid copy number to be approximately 24-27 copies per cell, as determined by the ratio of *bs2: flaA* DNA in *T. denticola* CF734. To confirm these results, we constructed *T. denticola* CF914 (Fig. 1A; Table 2; Supplemental Methods), in which the *bs2* gene is inserted in the *T. denticola* ATCC 35405 chromosome directly 3’ to *msp* (TDE0405) under transcriptional control of the *msp* promoter (P-*msp*). We obtained similar plasmid copy number results by qPCR comparing copies per cell of plasmid-encoded *bs2* in *T. denticola* CF734 versus chromosomally-encoded *bs2* in *T. denticola* CF914, using *flaA* as a normalization control between the two strains.

**Figure 1.**
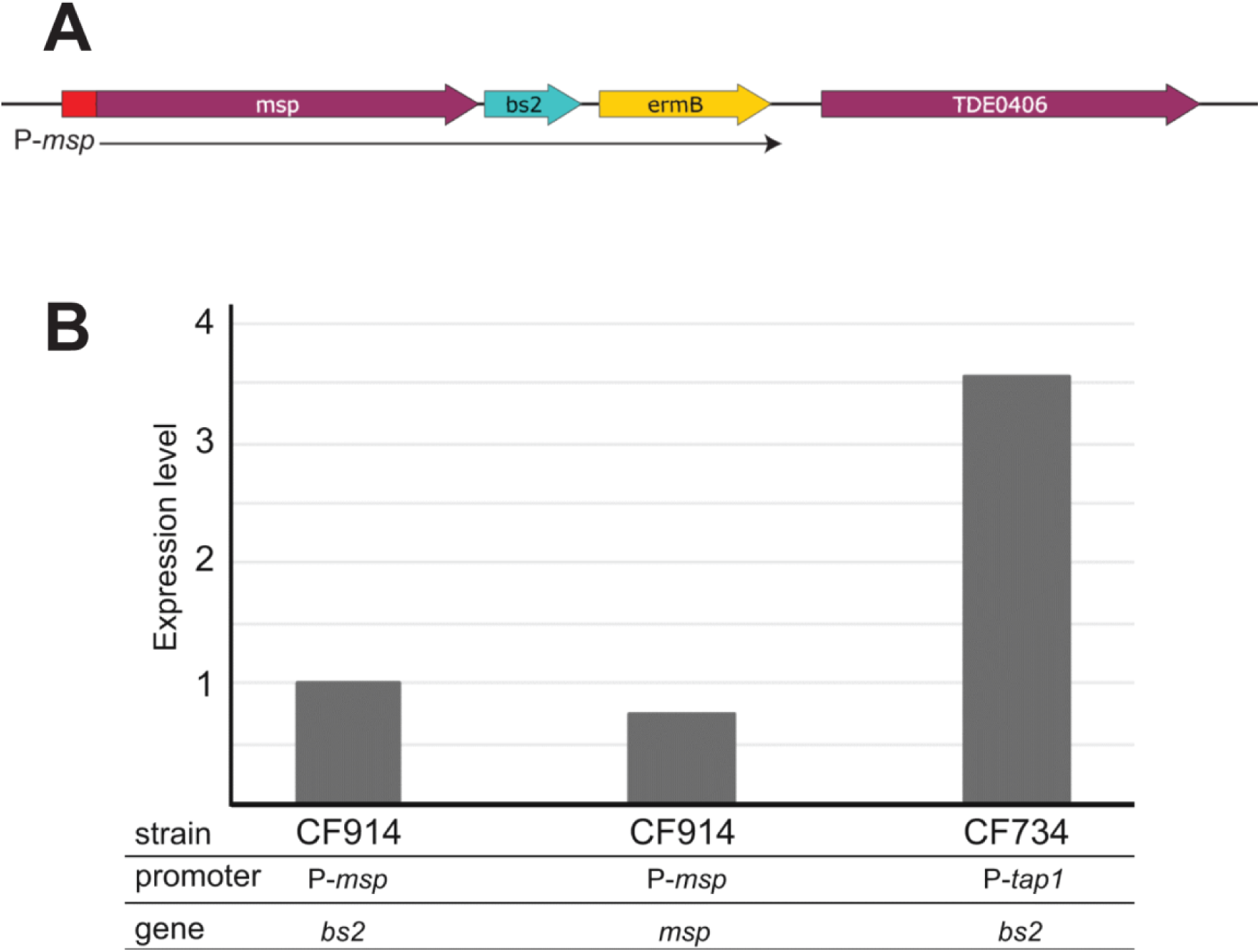
Transcription levels of *bs2* from *T. denticola* from a chromosomal locus and a plasmid. **Panel A:** Genetic construct for *bs2* expression from a *T. denticola* chromosomal locus. In *T. denticola* CF914, *bs2* and *ermB* are located directly 3’ to *msp.* As shown here, *msp, bs2,* and *ermB* are transcribed from the *msp* promoter (P-*msp*). **Panel B:** Relative transcription levels: *bs2* transcription in *T. denticola* CF914 is set at 1.0. Transcription of both *bs2* and *msp* localized on the chromosome of CF914 is driven by P-*msp.* Transcription of *bs2* on the shuttle plasmid in *T. denticola* CF734 is driven by P-*tap1*.

**Table 2.** *T. denticola* strains used in this study

| Strain | Plasmid | Features | Promoter<sup>a</sup> | Selection<sup>b</sup> | Source |
| --- | --- | --- | --- | --- | --- |
| 33520 | pTD1 | cryptic plasmid | N/A | N/A | ATCC |
| 35405 | none | wild type host for subsequent strains | N/A | N/A | ATCC |
| CF548 | none | $\Delta prcB$ mutant | *P-ermB* | Em | (24) |
| CF637 | none | $\Delta TDE0911$ (TdeIII) | N/A | Em | (18) |
| CF1090 | none | $\Delta TDE0911$; $\Delta msp$ | N/A | Em/Km | (46) |
| CF884 | pCF693 | pCF693 in ATCC 35405 | N/A | Thi | (29) |
| CF734 | pCF728 | *bs2* in pCF693 | *P-tap1* | Thi | (29) |
| CF914 | none | *bs2* on *T. denticola* chromosome | *P-msp* | Em | this study |
| CF1062 | pCF1056 | *bs2*-CO in pCF693Syn | *P-tap1* | Thi | this study |
| CF1086 | pCF1061 | *bs2*-CO in pCF693Syn | *P-msp* | Thi | this study |
| CF1179 | pCF1173 | *T. denticola* CF1090 host strain | *P-iTet* | Thi | this study |

<sup>a</sup> Promoter driving expression of cloned gene of interest. *P-tap1*: derived from *T. denticola* ATCC 33520 (21). *P-msp*: derived from *T. denticola* ATCC 35405. *P-ermB*: derived from pVA2198 (75).
<sup>b</sup> Antibiotic selection for chromosomal mutation or plasmid.

For comparative purposes, we found it useful to determine the copy number of pTD1. A prior study reported that pTD1 is present at 15-25 copies per cell, based on ethidium bromide staining of a plasmid dilution series (31). Using qPCR methodology with primers targeted to *flaA* and the pTD1 *rep* gene, we determined that cryptic plasmid pTD1 is present in *T. denticola* ATCC 33520 at approximately 9 copies per cell, somewhat lower than the previous report. We also determined that pTD1 has no potential as a component of a shuttle plasmid because its Rep protein is incompatible with the *E. coli* plasmid replicon (data not shown).

### Quantification of gene expression in *T. denticola* from diverse promoters

The original *E. coli-T. denticola* shuttle plasmids pKMR4 (17) and its derivatives pKMCou (19), pBFC (21) and pCF693 (29) differ in selectable markers and expression sites, and range in size between 6.4 kb and 8 kb. All carry the same 2.6 kb *T. denticola* replicon derived from pTS1, a 4.2 kb cryptic plasmid originally reported in an oral *Treponema* clinical isolate (20). Expression of genes cloned in plasmid vectors can have deleterious effects on host organisms, and this potential is amplified when using a two-host shuttle vector. This can be due to either (or both) the nature of the protein or to its expression level from the plasmid. Expression of genes cloned in shuttle vector pBFC and its derivatives is under the transcriptional control of P-*tap1*, a σ^28^-like promoter from the flagellar operon of *T. denticola* ATCC 33520 (37). This promoter was chosen because the flagellar genes are constitutively expressed in *T. denticola* at relatively high levels (38–40).

To investigate the effects of different promoters on expression of genes cloned in the shuttle plasmid, we used RT-qPCR to assess relative transcript levels of genes under control of several promoters of interest. To determine the relative activity of P-*tap1* in the shuttle plasmid, we compared *bs2* transcript levels in *T. denticola* CF914 and *T. denticola* CF734, where *bs2* is expressed from P-*msp* on the chromosome (Fig. 1A) or P-*tap1* on the shuttle plasmid (29), respectively. Using *flaA* as a normalization control (41), we determined that, as expected, transcription levels of *msp* and *bs2* in CF914 were similar (Fig. 1B). In contrast, *bs2* transcription was approximately 3.5 times greater in CF734 than in CF914. When combined with plasmid copy number determined above, we estimate that transcription is driven at levels approximately 7-fold higher by P-*msp* than by P-*tap1* when normalized to plasmid copy number. These results suggested that the *msp* promoter might be a candidate to drive enhanced expression of genes cloned in the shuttle plasmid.

We then assayed transcript levels of *fhbB* (TDE0108; encoding*T. denticola* FhbB (39)) and *ermB* (encoding a 23S rRNA methylase (42)). FhbB is an outer membrane Factor H-binding lipoprotein responsible for *T. denticola* resistance to complement-mediated killing (43), while ErmB is a well-characterized selectable marker used in *T. denticola* mutagenesis studies (16, 44). We hypothesized that genes encoding structural cell envelope-associated proteins would be more highly expressed than genes encoding cytoplasmic enzymes. In this case, a relatively weak promoter could be appropriate for recombinant expression of genes whose products are not well tolerated at high expression levels. Using isogenic *T. denticola* ATCC 35405-derived strains CF734, CF548 and CF914 (Table 2), we compared transcription levels of *ermB, msp*, *fhbB* and *Bs2*, using *flaA* as a normalization control throughout. As shown in Table 3, P-*ermB* was by far the weakest promoter, yielding less than one percent of the transcript levels of P-*fhbB* and P-*msp.* All of these promoters are on the chromosome of the relevant *T. denticola* strains tested.

**Table 3.**
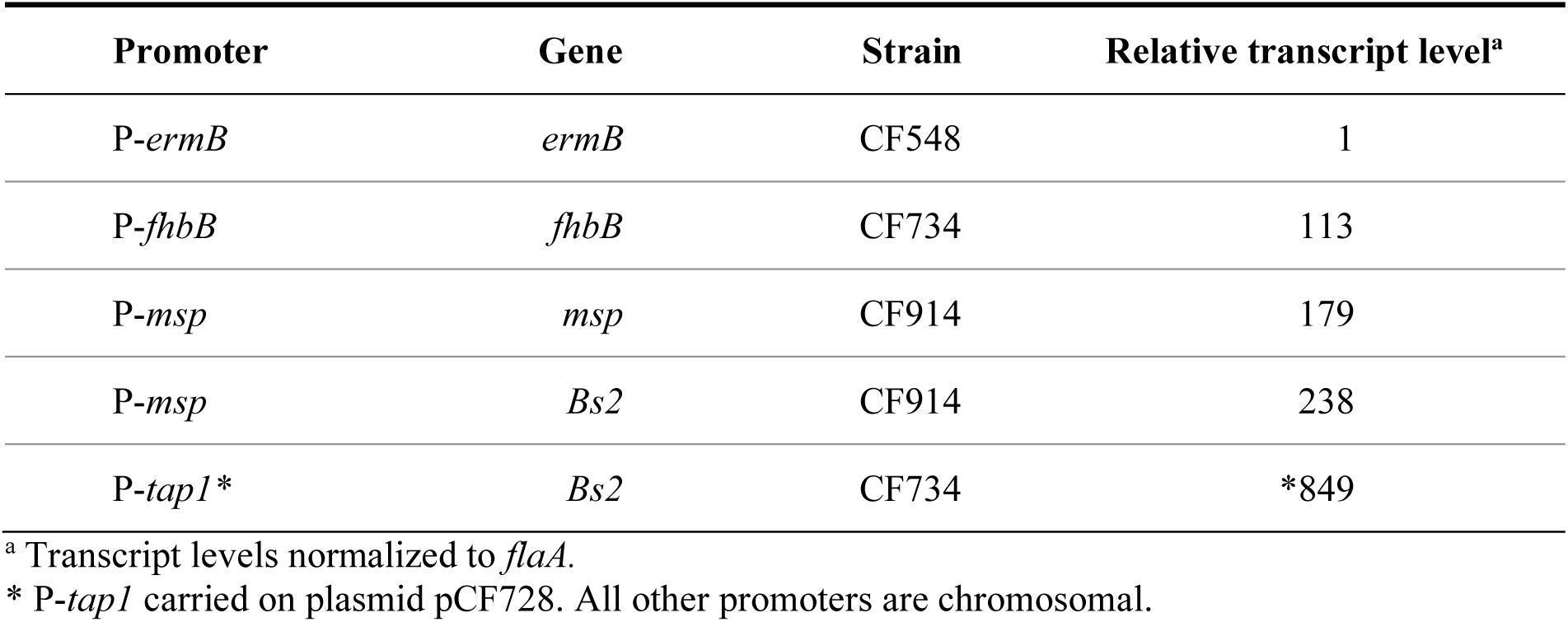
Relative transcription levels of promoter/gene pairs in *T. denticola*.

These results suggested P-*ermB* as an appropriate promoter for driving gene expression in the shuttle plasmid in instances where overexpression of a cloned gene might be problematic. For example, we have found that P-*ermB* is appropriate for use in *T. denticola* with genes encoding resistance to erythromycin (24), kanamycin (45), gentamicin (46) and chloramphenicol (unpublished data). We recently reported shuttle plasmid complementation of a *T. denticola* Δ*fhbB* mutant strain (43) using the shuttle plasmid carrying *fhbB* under transcriptional control of P-*ermB* (28). Given the shuttle plasmid copy number (24-27 per cell) and the relative activities of P-*fhbB* and P-*ermB* (Table 3), we estimate that FhbB expression would be approximately four times greater from the chromosome than from the shuttle plasmid. The complemented mutant expressed FhbB, though at lower levels than the parent strain, and the plasmid was stably maintained for multiple generations. In contrast, while a shuttle plasmid carrying *fhbB* under transcriptional control of P-*tap1* could be introduced into *T. denticola* Δ*fhbB*, it could not be stably maintained, presumably due to overexpression of FhbB (data not shown). Data in Table 3 suggests that expression of *fhbB* would be approximately eight times higher when driven by P-*tap1* on the shuttle plasmid than from its native promoter in the chromosome. This highlights the importance of rational pairing of promoters and genes of interest in shuttle plasmid systems for stable expression of cloned genes.

### Bs2 fluorescence in *T. denticola* as a reporter for promoter activity

The flavin mononucleotide-based fluorescent reporter protein Bs2 (36) is an attractive tool for studying gene regulation under anaerobic conditions, as has been previously documented in *Bacteroides* (47) and *Clostridium* (48). We utilized several plasmid and chromosomal constructs to demonstrate the utility of Bs2 fluorescence to monitor promoter activity in *T. denticola*. As noted above, *T. denticola* CF914 carries the *bs2* gene directly 3’ to *msp*, under transcriptional control of the *msp* promoter (P-*msp*; Fig. 1A). The *Bs2* gene carried in *T. denticola* CF914 (Table 2) and on shuttle plasmid pCF728 (Table 1) are optimized for expression in *Clostridium spp.* (49), which have genomic G-C contents similar to that of *T. denticola.* As shown in Fig. 2, *bs2* specifically codon-optimized for *T. denticola* (*bs2-*CO) differs in 54 of 366 nucleotides from the *bs2* coding sequence used in pCF728 (29). For cloning purposes, an additional conservative base change (A^225^→T) was made to remove an internal PstI site. The *bs2-*CO gene was cloned in pCF693Syn such that its transcription in the resulting plasmid (pCF1056; Fig. 3) is driven by P-*tap1*, exactly as in pCF728. To maximize *bs2-*CO transcription in *T. denticola* we also replaced P-*tap1* with P-*msp*, yielding pCF1061 (Fig. 3). Consistent with our prior observation that the *tap1* promoter is not active in *E. coli* (29), Bs2-dependent fluorescence was expressed in *E. coli* from pCF1061 but not from pCF1056 (Fig. 4). Each of these plasmids was then introduced into *T. denticola* ATCC 35405, yielding *T. denticola* strains CF1062 (ATCC 35405/pCF1056) and CF1086 (ATCC 35405/pCF1061) (Tables 1, 2).

**Figure 2.**
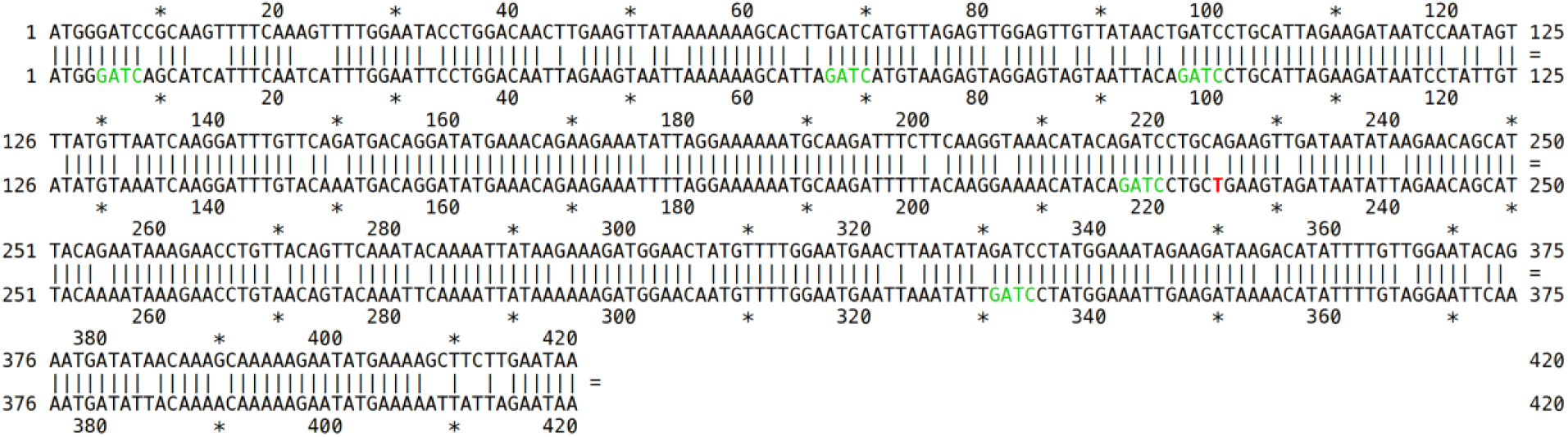
Codon optimization of *bs2* for expression in *T. denticola.* Codon-optimized *bs2* is shown below the starting *bs2* DNA sequence from pCF693. A conservative single base change at base 220 (A→T, red in *bs2-*CO) was made to alter a PstI site (221–226). Five recognition sites for TdeI (green in *bs2-*CO) were not removed.

**Figure 3.**
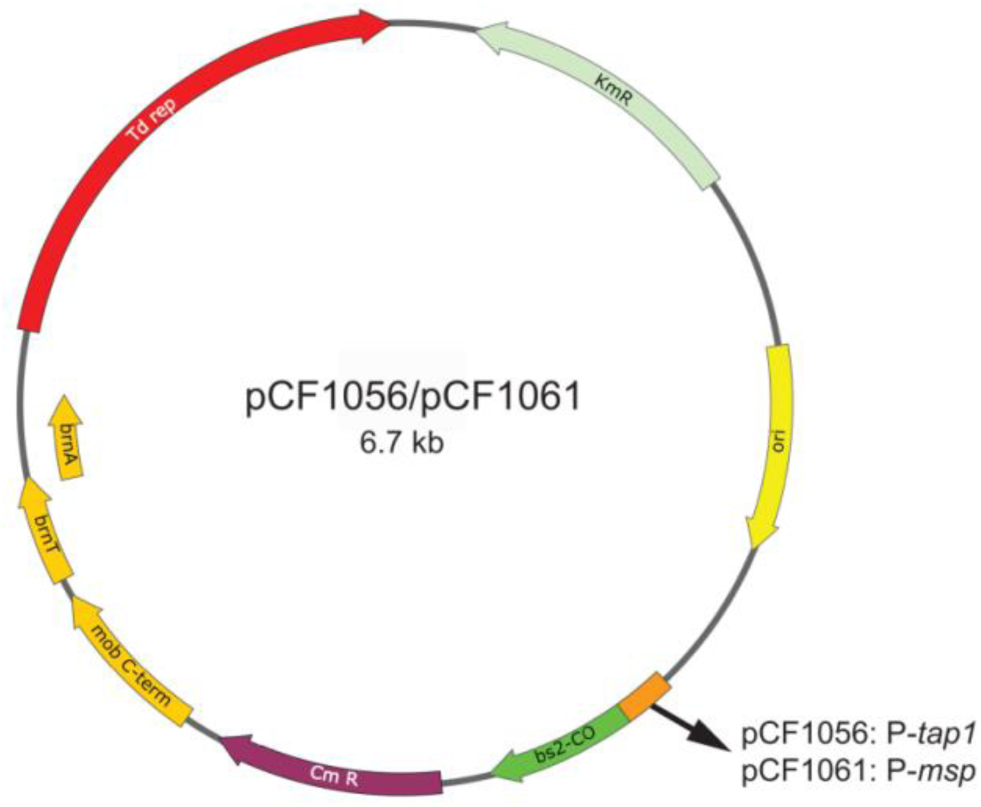
Plasmids for expression of *bs2*-CO in pCF693Syn. Both pCF1056 and pCF1061 carry the *bs2*-CO gene in pCF693Syn. Transcription of *bs2*-CO is driven by either P-*tap1* (pCF1056) or P-*msp* (pCF1061).

**Figure 4.**
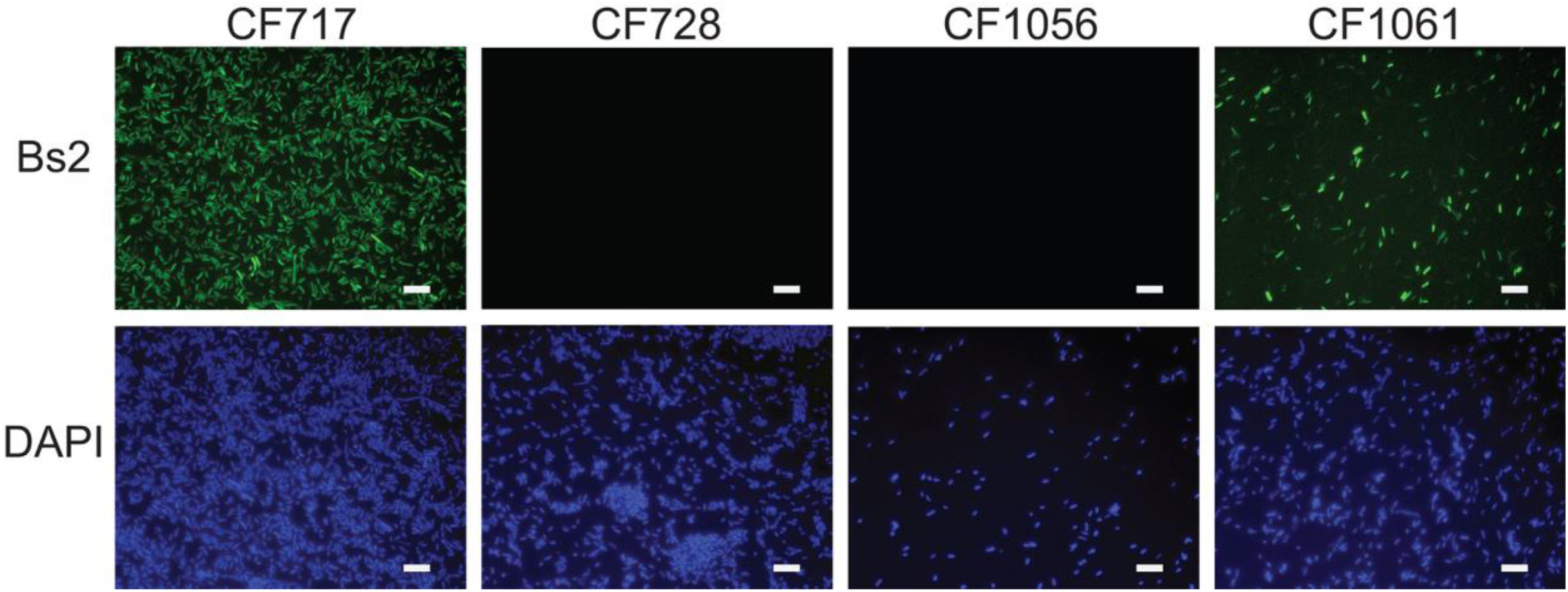
Bs2 fluorescence expression in *E. coli* from different promoters. Expression of Bs2 from plasmid vectors in *E. coli* (Table 1) under transcriptional control of a *Clostridium* promoter in pCF717 (CF717), *T. denticola* P-*tap1* promoter in pCF728 and pCF1056 (CF728, CF1056), and the *T. denticola* P-*msp* promoter (CF1061). Scale bar: approximately 5 µm.

Fluorescence expression in *T. denticola* strains carrying the *bs2-*CO gene was monitored spectrophotometrically at optimal emission and excitation settings (Ex:449 nm; Em: 495 nm) and is reported in Table 4. Codon optimization of *bs2* resulted in a modest increase in fluorescence expression in *T. denticola* (data not shown). Replacing promoter P-*tap1* with P-*msp* resulted in nearly three-fold higher fluorescence expression in *T. denticola* CF1086 compared with *T. denticola* CF1062 (Table 4). Fluorescence values in *T. denticola* carrying pCF1061 were much lower than those observed in *E. coli* carrying the same plasmid (data not shown). This is likely due to the fact that the *E. coli* replicon carried on the shuttle plasmid is derived from the high copy number vector pUC19 (50), while the copy number of the shuttle plasmid in *T. denticola* is approximately an order of magnitude lower due to its *Treponema-*specific replication mechanism (51).

**Table 4.** Bs2 fluorescence expression in *T. denticola*

| Strain name | *T. denticola* / plasmid | Features | Mean RFU/OD<sub>600</sub><sup>a</sup> | Std Error |
| --- | --- | --- | --- | --- |
| CF1060 | 35405 / pCF693Syn | empty vector | 0.306 | 0.097 |
| CF1062 | 35405 / pCF1056 | *Ptap1-bs2-CO* | 0.596 | 0.040 |
| CF1086 | 35405 / pCF1061 | *Pmsp-bs2-CO* | 1.64 | 0.195 |

<sup>a</sup> RFU/OD<sub>600</sub>: Relative fluorescence units/OD<sub>600</sub>; excitation 449 nm; emission 495 nm (5 nm bandwidth). Data shown are from three biological replicates with triplicate samples.

As shown in Table 4, we detected considerable intrinsic autofluorescence in *T. denticola* ATCC 35405. We conducted a separate series of experiments in which minor variations were included in attempts to minimize autofluorescence. Growth media used for *T. denticola* growth media contains yeast extract and serum, both of which exhibit strong fluorescence in the 500 nm range (52). However, neither increasing the numbers of PBS washes nor reducing the excitation and emission bandwidths from 12 nm to 5 nm decreased *T. denticola* autofluorescence (data not shown). Similarly, based on the complete spectral range of emission/excitation data from *T. denticola* ATCC 35405, we tested an array of combinations of excitation and emission settings that could potentially reduce autofluorescence while retaining sensitive detection of Bs2 fluorescence. While the results showed some very minor differences, we were unsuccessful in significantly reducing *T. denticola* autofluorescence in this assay (data not shown).

We focused on quantitative analysis of Bs2 fluorescence for two reasons. Firstly, while Bs2 has been utilized by us and others in fluorescence microscopy of oral microbes (29, 53), its primary utility is in biofilm studies (54) and as a quantitative reporter of promoter activity (47, 54–56). Secondly, compared with microscopy, spectophotometric assays are rapid and yield consistent, quantitative results. Several factors contribute to technical challenge of detecting Bs2 expression in *T. denticola* by fluorescence microscopy. These include its relatively unfavorable quantum yield and extinction coefficient, intrinsic autofluorescence of *T. denticola* compared with *E. coli,* and rapid photobleaching during the relatively long times required for epifluorescence image acquisition (54). This last issue was also noted in studies of *Veillonella* (54) and *Clostridium* (57). For this reason, we anticipate that the most promising use of Bs2 in *T. denticola* will be in monitoring activity of promoters of interest, as described here.

### Tet-inducible promoter system for calibrated gene expression in *T. denticola*

Complementation of mutant strains provides an important level of rigor to mutagenesis experiments. Historically, few studies of *T. denticola* mutant strains have included this component, due to limitations of earlier versions of the shuttle plasmid system and, until quite recently, the small number of validated selectable markers. Other than complementation of flagellar component mutants in *T. denticola* ATCC 33520 (17, 19, 21) using a shuttle plasmid, all other complementation studies in *T. denticola* have been done by insertion of the complementing gene in the mutant chromosome, either in *cis* (25, 41) or in *trans* (26).

Msp, an oligomeric β-barrel outer membrane protein (58) present at approximately 2×10^5^ copies per cell (59), is an ortholog of the *T. pallidum* Tpr gene family that includes TprK, whose remarkable *in vivo* hypervariability is proposed to contribute to *T. pallidum* immune evasion (60). Attempts at complementation of *T. denticola* Δ*msp* mutant strains has proved problematic due to the properties of the Msp protein. The primary issue is that expression of full-length Msp including its signal peptide is toxic in *E. coli*, presumably due to disruption of membrane integrity (61). Because we had previously observed that the P-*tap1* promoter is not active in *E. coli* ((29) and Fig. S2), we hypothesized that we could clone *msp* in pCF693 under control of P-*tap1* for use in complementation experiments. We first constructed pCF1112, which carries the full length *msp* gene with its native terminator in pCF693Syn under transcriptional control of P-*tap1* (Table 1). Because it has never been determined whether the predicted promoter sequence immediately 5’ to the Cm^R^ gene in pCF693 is active in *T. denticola*, we also constructed pCF1121 in which the *msp* transcription terminator sequence in pCF1112 is deleted, so that transcription would proceed from *msp* into the gene encoding CmR. We were unable to recover transformants of *T. denticola* Δ*msp* strain CF1090 carrying either of these plasmids, suggesting that the issue was with Msp expression rather than with the selectable marker. Based on analysis of relative promoter activity, we anticipated that *msp* transcription from P-*tap1* on the shuttle plasmid in *T. denticola* would be 3-4 fold higher than the already very high level in the wild type parent strain (Fig. 1 and Table 3). This suggested that overexpression of Msp driven by P-*tap1* on the shuttle plasmid was not tolerated by *T. denticola*.

To expand the range of expression control in the shuttle plasmid, we then constructed pCF1173 in which expression of Msp is under control of a tetracycline-inducible promoter cassette (*P-tet*) that contains the *tetR* gene and its constitutive promoter and (in the opposite orientation) a promoter interrupted by a TetR-binding operator sequence (*tetO*) such that, in the absence of a tetracycline analog (anhydrotetracycline; ATc), transcription of the gene of interest is blocked by TetR binding to *tetO* (62, 63). The *msp* coding region was fused to the *P-tet* promoter cassette and cloned in pCF693Syn, resulting in pCF1173 (Fig. Fig. 5A, 5B; Table 1). This plasmid was introduced into *T. denticola* Δ*msp* strain CF1090 (Table 1). As shown in Fig. 5C and 5D, calibrated expression of Msp was induced by addition of increasing amounts of ATc to the growth medium. Importantly, the complemented strain *T. denticola* CF1179 (Table 2) expressed both monomeric and oligomeric forms of Msp, as is also seen in the wild type strain (Fig. 5C). Importantly, Msp protein levels in the complemented mutant CF1179 increased as cells were grown in increasing levels of ATc. However, we were not able to attain wild type levels of Msp expression from pCF1173. This may be because the P-*xyl/tetO* promoter (derived from *Staphylococcus* (64) in the Tet-inducible cassette is weaker than the native *T. denticola* promoter. Alternatively, it is possible that ATc loses effective activity over the *T. denticola* growth phase due to its relatively short half-life under culture conditions (65). Studies are in progress to further optimize this system for *T. denticola* and other slow-growing anaerobes.

**Figure 5.**
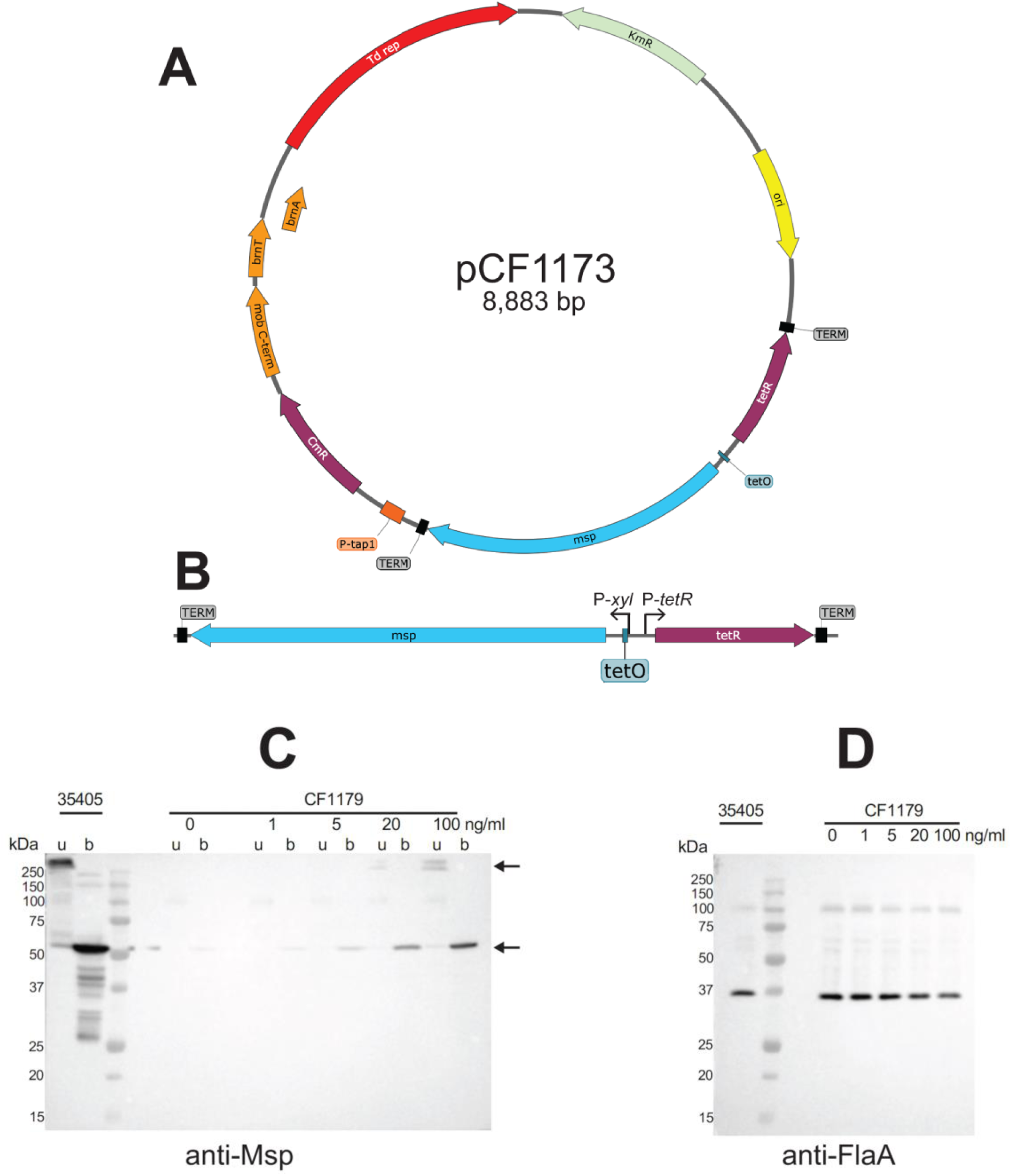
Inducible expresson of Msp in a complemented *T. denticola* Δ*msp* mutant. *T. denticola* CF1090 (35405 Δ*msp*) was transformed with shuttle plasmid pCF1173 (Panel A), which carries *msp* under transcriptional control of the tetracyline-inducible promoter derived from pRPF185 (62) (Panel B), yielding complemented Δ*msp* mutant strain CF1179. Expression of Msp in *T. denticola* parent ATCC 35405 and complemented Δ*msp* mutant CF1179 was assayed by Western immunoblots probed with antibodies raised against native Msp (Panel C) and FlaA (Panel D). CF1179 was grown in concentrations of ATc ranging from 0 to 100 ng/mL to induce Msp expression. Samples probed with anti-Msp were held on ice (**u**) or boiled (**b**) prior to electrophoresis to detect Msp in monomers and oligomers (arrows), respectively.

### Summary

We provide here expanded characterization of a *T. denticola* shuttle plasmid and demonstrations of its expanded use in expression and complementation studies in this periodontal pathogen. In addition to new information on the plasmid components, we demonstrated its utility for genetic complementation of isogenic mutant strains and as a vector for spectrophotometric quantitation of promoter activity and expression of an oxygen-independent fluorescent protein. Finally, we report the first application of the powerful Tet-inducible calibrated promoter system in *Treponema.* These results will facilitate expanded genetic studies in *Treponema* species that are required to understand their roles in interactions with other microbe and with host cells and tissue.

## MATERIALS AND METHODS

### Bacterial strains and growth conditions

*T. denticola* strains (Table 2) were grown as previously described under anaerobic conditions in TYGVS broth or semisolid agar medium (14) supplemented with erythromycin (Em; 40 µg ml^-1^), kanamycin (Km; 25 µg ml^-1^) or thiamphenicol (Thi; 10 µg ml^-1^) as appropriate. We used the chloramphenicol analog thiamphenicol because it is more stable over the long incubation period required to grow *T. denticola* (66). Unless otherwise noted, growth media contained 5% rabbit or horse serum. All *T. denticola* growth media were incubated under anaerobic conditions for at least 18 h prior to use (67). Purity of spirochete cultures was monitored by darkfield microscopy. *E. coli* strains JM109 and JM110 (50), used for plasmid construction and propagation, respectively, were grown in LB medium (68) supplemented as appropriate with carbenicillin (Cb; 50 µg ml^-1^), kanamycin (Km; 50 µg ml^-1^).

### Construction of recombinant plasmids

Shuttle plasmids used to transform *T. denticola* were based on either pCF693 (29) or pCF693Syn (28). Plasmid pCF693Syn was designed to be resistant to all *T. denticola* ATCC 35405 R–M systems, while conserving amino acid sequences of plasmid-encoded genes in pCF693 and the nucleotide sequence of the P-*tap1* promoter that drives expression of genes cloned in its expression site. Plasmids were constructed using either restriction fragments of previously verified plasmids or PCR products generated with high-fidelity DNA polymerases (Phusion (New England Biolabs, Beverly, MA) or Taq-HF (Invitrogen, Carlsbad, CA)) using oligonucleotide primers listed in Table S1. Details of individual plasmid constructs are found in Supplemental Material. Plasmid constructs were isolated after transformation into *E. coli* JM109. Plasmids used to transform *T. denticola* were propagated in *E. coli* JM110 (*dam^-^, dcm^-^*). Recombinant plasmid constructs isolated from *E. coli* before and after passage through *T. denticola* were verified by DNA sequencing at the University of Michigan DNA Sequencing Core Facility or by Eurofins (Eurofins Genomics, Louisville, KY) and analyzed using DNASTAR sequence analysis software (DNASTAR Inc., Madison, WI).

### Transformation of *T. denticola*

Preparation and electroporation of competent *T. denticola* cells were conducted as previously described (15, 29). For allelic replacement mutagenesis, *T. denticola* competent cells were electroporated with linear double stranded DNA consisting of a selectable marker gene cassette flanked by *T. denticola* DNA of the target region. Details of chromosomal insertion of *bs2* 3’ to the *msp* gene (TDE0405) are shown in Supplemental Methods. For plasmid transformations, *T. denticola* competent cells were electroporated with unmethylated shuttle plasmid DNA that had been propagated in *E. coli* JM110. Following electroporation, cells were incubated in non-selective medium for 18h, then plated in pre-reduced TYGVS agar medium containing thiamphenicol for selection of transformants. Presence of a shuttle plasmid in transformed *T. denticola* was verified in broth cultures grown from individual colonies. Standard polymerase chain reaction (PCR) using Platinum Taq polymerase (Life Technologies) and shuttle plasmid-specific oligonucleotide primers CX930 and CX931 (Table S1) were used to amplify a 2.5 kb region of the shuttle plasmid including the 3’ end of the Cm^R^ gene through the middle of the pTS1-derived *rep* gene. For further validation, plasmid DNA isolated from Thi^R^ *T. denticola* cultures was transformed back into *E. coli.* The restriction enzyme digestion pattern of plasmids subsequently isolated from Km^R^ *E. coli* transformants was compared with that of original shuttle plasmid and the DNA sequence of either the cloned insert or the entire plasmid was validated.

### Codon optimization

The gene encoding the Bs2 fluorescent protein was modified for optimized expression in *T. denticola* using the JCat codon usage optimization tool (69) and synthesized by GeneArt (ThermoFisher Scientific). The synthesized gene (*bs2-*CO) lacks recognition sites for *T. denticola* restriction-modification enzymes other than TdeI (G^m6^ATC; REBASE, http://rebase.neb.com). Additional modifications were removal of an internal PstI site and addition of SpeI and PstI sites at the 5’- and 3’-ends of the coding region, respectively, to facilitate cloning in pCF693Syn.

### Promoter optimization

To test effects of different *T. denticola* promoters on Bs2 expression, *bs2-*CO was cloned between the SpeI and PstI sites of pCF693Syn such that its transcription in the resulting plasmid (pCF1056) was directed by the *T. denticola tap1* promoter (P-*tap1*) on the plasmid vector, as in pCF728 (29). To replace P-*tap1* in pCF1056 with P-*msp,* a 149-bp fragment directly 5’ to the *T. denticola msp* coding region was amplified and cloned in pSTBlue-1. The resulting plasmid (pCF540) was then used as one template for ligation-independent PCR cloning (70), while pCF1056 was used as the second template, using oligonucleotide primers containing overlapping P-*msp*/pCF1056 sequences (Table S1). Following the amplification reaction, the products were combined, digested with DpnI to remove methylated plasmid template and used to transform *E. coli* JM109 (70). In the resulting plasmid pCF1061, transcription of *bs2-*CO is driven by the *msp* promoter.

### Quantitative PCR

Plasmid copy number was determined using quantitative PCR (qPCR) from plasmid-containing *T. denticola* strains (71). Fresh broth cultures (200 µl) were pelleted and resuspended in 100 µl distilled water, from which 1 µl was used as template for qPCR. For thermal cycling, primers designed to amplify 80-100 bp from a chromosomal gene (*flaA*) and a plasmid gene (*bs2* or pTD1 *rep,* depending on plasmid type; Table S1) were used with QuantiTect™ SYBR Green PCR Kit (Qiagen; https://www.qiagen.com/us/) in 25 µl of reaction buffer in a MyiQ™ Single-Color Real-Time PCR Detection System (Bio-Rad Laboratories, Hercules, CA). Reaction conditions were 95°C for 15 min, followed by 40 cycles of 95°C for 15 sec, 60°C for 60 sec, and 68°C for 30 sec. Each individual assay was performed in triplicate. Threshold values were calculated using baseline cycles 2 to 10. Data were analyzed using by the ^ΔΔ^C_T_ method (72) using software supplied by the manufacturer.

### Quantitative reverse transcriptase PCR (RT-qPCR)

Total RNA was extracted from *T. denticola* cultures, treated with DNAse and reverse transcribed to cDNA using random hexamer primers as described previously (41). Gene-specific primers were designed using the NCBI-PrimerBlast Tool (https://www.ncbi.nlm.nih.gov/tools/primer-blast/) to have similar annealing temperatures and amplicon sizes of 80-100 bp. (Table S1). One µL of the resulting cDNA was amplified using the QuantiTect™ SYBR Green PCR Kit (Qiagen; https://www.qiagen.com/us/) in 25 µl of reaction buffer, with *T. denticola flaA* serving as an internal reference control for normalization between samples. Thermal cycling was performed in a MyiQ™ Single-Color Real-Time PCR Detection System (Bio-Rad Laboratories, Hercules, CA) at 95°C for 15 min, followed by 40 cycles of 95°C for 15 sec, 60°C for 60 sec, and 68°C for 30 sec. Each individual assay was performed in triplicate. Threshold values were calculated using baseline cycles 2 to 10. Data were analyzed using by the ^ΔΔ^C_T_ method (72) using software supplied by the manufacturer. Experiments were performed on three biologically independent replicates, each containing triplicate samples. Transcription data are presented as means of triplicate samples.

### Tetracycline induction assay in complemented *T. denticola* mutant

*T. denticola* CF1179 grown in TYGVS medium containing thiamphenicol (Thi^10^) was used to inoculate separate cultures in the same medium supplemented with anhydrotetracyline (ATc; Cayman Chemical, Ann Arbor, MI) at final concentrations of 0, 1, 5, 20 and 100 ng/mL. ATc was added to actively growing cultures, and *T. denticola* cells were harvested for analysis after 24 h growth.

### Protein electrophoresis and immunoblotting

Protein gel electrophoresis and western immunoblotting were carried out as described previously (Fenno et al., 1996). First, the optical density 0.2 at 600 nm (OD_600_) of *T. denticola* cultures was determined, then cells were harvested by centrifugation at 10,000 g (10 min, 4°C) in the presence of 2 mM phenylmethylsulfonyl fluoride (PMSF), washed once in an equal volume of PBS containing 2 mM PMSF, then suspended in a volume of standard SDS–PAGE sample buffer containing dithiothreitol and 2 mM PMSF equal to 0.5x the starting OD_600_. Prior to electrophoresis, cells were lysed and genomic DNA was sheared by multiple passages through a 26-guage needle. Samples were heated at 100°C for 5 min or not heated prior to electrophoresis, and equal volumes of cell lysate samples were loaded on gels. Proteins blotted from gels to nitrocellulose membranes were detected with polyclonal antibodies raised in in rabbits against FlaA (73) or native Msp (74), followed by appropriate horseradish peroxidase (HRP)-conjugated goat anti-rat or anti-rabbit IgG (Thermo Scientific, Rockford, IL). Protein bands of interest were visualized using SuperSignal West Pico chemiluminescent substrate (Thermo Scientific).

### Epifluorescence detection of Bs2 in *E. coli* cells

*E. coli* strains were grown in LB broth with appropriate antibiotics. One mL samples of overnight cultures were washed once in PBS, then resuspended in one mL PBS. Approximately 10 µL of cell suspension was spread on each slide and allowed to air-dry. Attached cells were fixed with ice-cold 100% methanol for 2 min, rinsed with water and air-dried. Coverslips were mounted with Pro-Long Gold antifade reagent containing DAPI (Invitrogen/Thermo Fisher P36935). Slides were analyzed on an Olympus BX40 microscope under 600x magnification fitted with standard filter sets: blue (excitation/emission: 375/450 nm); and green (excitation/emission: 485/535 nm). Images were acquired with an Olympus-DP73 camera using Olympus cellsSens software.

### Spectrophotometric detection of Bs2 fluorescence

Broth cultures of *T. denticola* (three-day) were washed twice in PBS and suspended in PBS. Samples (0.2 ml) were loaded in 96-well glass bottom plates (Cellvis, Mountain View, CA). Using a Varioskan Flash instrument (ThermoScientific, Vantaa, Finland) according to the manufacturer’s instructions, we recorded both OD_600_ and fluorescence (excitation at 449 nm, emission 495 nm; 5 nm bandwidth; 100 ms). Results are expressed as relative fluorescence units (RFU) divided by OD_600_.

## Supporting information

Supplemental Methods, Table, DNA sequence

## ACKNOWLEDGEMENTS

We dedicate this work to the memory of Dr. Howard Kuramitsu, a pioneer in *Treponema* genetic analysis. This work was supported by NIH grants DE025225 and AI105362 (J.C.F.). P.S. was supported by the Department of Biologic and Materials Sciences and Prosthodontics, University of Michigan School of Dentistry. B.R.C. and S.M.A. were supported by the University of Michigan Undergraduate Research Opportunity Program.

## Notes

### Competing Interest Statement

The authors have declared no competing interest.

