## Supplemental Methods, Table, DNA sequence for "Functional characterization and optimization of protein expression in *Treponema denticola* shuttle plasmids"

### SUPPLEMENTAL MATERIAL

#### SUPPLEMENTAL METHODS

**Construction of *T. denticola* CF914 carrying the *bs2* gene in the chromosome.** To construct a *T. denticola* strain expressing Bs2 from the *T. denticola* chromosome, we first constructed an *E. coli* plasmid (pCF639) by overlap extension PCR cloning (1) carrying the following 3.25 kb insert: 3' end of *msp* (TDE0405); *ermB* gene encoding $Em^R$ (with its ribosome binding site); 5' end of TDE0406. We then amplified the *bs2* gene with its ribosome binding site from pCF728 (2) using primers CX1100 and CX1101 (Table S1)) and cloned it into the XhoI site directly 3' to the *msp* stop codon in pCF639. The resulting plasmid (pCF862) was digested in PvuII to release the vector sequence and used to transform *T. denticola* by electroporation. Transformants were selected for $Em^R$, as described previously (3). As illustrated in Fig. 1A, when this construct is inserted into the *T. denticola* chromosome by double crossover homologous recombination, transcription of both *bs2* and *ermB* in the resulting *T. denticola* CF914 are driven by the *msp* promoter (4).

**Plasmid construction for complementation of *T. denticola* Δ*msp* mutant.** The coding region of *msp* was PCR-amplified from ATCC 35405 genomic DNA using oligonucleotide primers CX1630 and CX1631 (Table S1) and cloned by overlap extension PCR with pCF693 DNA that had been linearized by PCR-amplification using oligonucleotide primers CX1632 and CX1633 (Table S1). In the resulting plasmid (pCF1112; Table 1) the full length *msp* gene and terminator signal are directly 3' to vector-encoded P-*tap1.* Two subsequent derivative plasmids were made from pCF1112. The *msp* terminator signal was removed by amplifying pCF1112 with oligonucleotide primers CX1644 and CX1645, which flank the terminator site and contain BglII sites. The amplicon was digested in BglII and self-ligated to yield pCF1121.

To modulate Msp expression in a smaller version of the shuttle plasmid, we first replaced the segment of pCF693 encompassing P-*tap1* and the $Cm^R$ gene with the *msp* gene under transcriptional control of either P-*tap1* or an inducible tetracycline promoter (P-*tet*; (5)). The inducible P-*tet* expression cassette from pRPF185 (6) was PCR-amplified with primers CX1687 and CX1688, which contain 5'-tails complementary to pCF693 (residues 2378-2395) and *msp* (residues 1-22), respectively. The *msp* coding region and terminator sequence were amplified from *T. denticola* DNA with primers CX1689 and CX1690, which contain 5'-tails complementary to P-*tet* (pRPF185 residues 4026-4042) and pCF693 (residues 3488-3503). The fragments were joined by ligation independent cloning (7) then transformed into *E. coli* JM109. Following validation of the resulting plasmid pCF1173 DNA sequence, the plasmid was propagated in *E. coli* JM110 to yield unmethylated plasmid DNA, which was then introduced into *T. denticola* CF1099 by electroporation as described previously (2). Colonies carrying the plasmid were selected by resistance to thiamphenicol. Expression of *msp* in strains carrying the inducible P-*tet,* was induced by addition of anhydrotetracycline (1-100 ng/mL) to actively growing cultures. Expression of Msp was monitored by immunoblot with anti-Msp and anti-FlaA antibodies. Complemented strains were validated by recovery of the plasmid, which was confirmed by DNA sequencing following reintroduction into *E. coli.*

**TABLE S1. Oligonucleotide primers used in this study.**

| Primer | Sequence | Target |
|---|---|---|
| CX233 | GCAAAGATAGCAAACTTTATCC | *ermF/B* 5' end (F) |
| CX236 | AGCGACTCATAGAATTATTTCC | *ermF/B* 3' end (R) |
| CX661[a] | GGGCGGCCGCTTTGTTGTTAATATGCCG | NotI-P-*msp* (F) |
| CX662[a] | CCATGGAAAAAATTCCTCCTTGTTATTTG | NcoI-P-*msp* (R) |
| CX930 | CGGCCAAGTTCTATACGCTGATGA | pTS1 *rep* (R) |
| CX931 | ATTGTTTCCCAAAACACCTATACCTGA | pBFC *cmR* (F) |
| CX1100[a] | TATTATCTCGAGAGGAGGTTCACTAGTA | XhoI-RBS-*bs2* (F) |
| CX1101[a] | TATCATCTCGAGTTATTCAAGAAGCTTTTCA | XhoI- *bs2* (R) |
| CX1196 | TGACGGCTGGAGAGAATTGG | *flaA1*-qPCR (F) |
| CX1197 | GGATAAAGCCTCAATTCCCTAGACT | *flaA1*-qPCR (R) |
| CX1206 | GGAGCTGCTTTTTACGTGGT | pTS1 *rep*-qPCR (F) |
| CX1207 | GTAGGCAAGCGCTGATAGTT | pTS1 *rep*- qPCR (R) |
| CX1208 | GCGACCGATTACATCAAGGA | pTD1 *rep*-qPCR (F) |
| CX1209 | CCATAAGCCAATAAACGGCG | pTD1 *rep*-qPCR (R) |
| CX1210 | AGGATTTGTTCAGATGACAGGA | *bs2*-qPCR (F) |
| CX1211 | ACTTCTGCAGGATCTGTATGT | *bs2*-qPCR (R) |
| CX1230 | TATAAACGCGCTTGAGAATG | *fhbB*-qPCR (F) |
| CX1231 | AATGCAAGGGCTTCAGTATT | *fhbB*-qPCR (R) |
| CX1232 | TAAAACTTACCCGCCATACC | *ermB*-qPCR (F) |
| CX1233 | CGATATTCTCGATTGACCCA | *ermB*-qPCR (R) |
| CX1259 | CAGCTTCAGGAGATACGAATTTTGTTG | *msp*-qPCR (F) |
| CX1260 | TGGGAACTGCGCTTAAGAGATGG | *msp*-qPCR (R) |
| CX1591 | ATGGGATCAGCATCATTTCAATC | *bs2* internal (F) |
| CX1592 | TCAGCAGGATCTGTATGTTTTCCT | *bs2* internal (R) |
| CX1572[b] | TCAAGCTTGGTACTTTGTTGTTAATATGCCGAAAAAAG | P-*msp* / *bs2* (F) |

| | | |
|---|---|---|
| CX1573[b] | ATGATGCTGATCCCATAAAAAATTCCTCCTTGTTATTTG | P-*msp / bs2* (R) |
| CX1574[b] | TAACAAGGAGGAATTTTTTATGGGATCAGCATCATTTC | *bs2* / pCF1056 (F) |
| CX1575[c] | GCATATTAACAACAAAGTACCAAGCTTGATGCAATC | P-*msp /* pCF1056 (R) |
| CX1590[d,e] | ATGATGATGATGATGGCTGCTGCCCATAAAAAATTCCTCCTTGTTATTTGT | 6xHis-*P-msp* (R) |
| CX1630[f] | GATCAATGTATAGGAGGTTCTTTTATGAAAAAAATTCTGG | pCF693-5'end *msp* (F) |
| CX1631[f] | GATGGATATCTGCAGGAGAATAGCAGCAGAG | *msp* T-pCF693 (R) |
| CX1632[f] | TAGACTCTGCTGCTATTCTCCTGCAGATATCCATCACAC | pCF693- *msp* T (F) |
| CX1633 | CGCCAGAATTTTTTCATAAAAGAACCTCCTATACATTGATCCTA | 5'end *msp-* pCF693 (R) |
| CX1638 | TATGTCGACAGGAAACAGCTATGACCATGA | Δ*lacPO* in pCF693 (F) |
| CX1639 | ATTGTCGACATTGCGTTGCGCTCACTG | Δ*lacPO* in pCF693 (R) |
| CX1640 | TATAGATCTTGTGTCTAATAGACTCTGCTG | Δ*msp*T (F) |
| CX1641 | ATTAGATCTCGGCGGGTTTTAAACTATAC | Δ*msp*T (R) |
| CX1644[a] | TATAGATCTAGACTCTGCTGCTATTCTCC | Δ*msp*T in pCF1112-F |
| CX1645[a] | ATTAGATCTTAAACTATACTAGGTATAGATTAGTAGATAA | Δ*msp*T in pCF1112-R |
| CX1687[a] | CAACGCAATGTCGACCTAATTTTTAGACTTAAGGG | P1: 693.SalI-Ptet-F |
| CX1688[f] | AAATCGCCAGAATTTTTTTCATAAAATTTTCTCCTTTACT | P2: Ptet-msp-R |
| CX1689[f] | GTAAAGGAGAAAATTTTATGAAAAAAATTCTGGCGATTTT | P3: Ptet-msp-F |
| CX1690[a] | TCGGCATTATCTCATAGTCGACGAGAATAGCAGCAGAGTCTA | P4: msp-SalI-pCF693-R |
| CX169 [a] | GCAATGTCGACCTAATTTTTAGACTTAAGGG | P1x:OEPCR P1/2+P3/4 |
| CX1692[a] | ATAGTCGACGAGAATAGCAGCAGAGTCTA | P4x:OEPCR P1/2+P3/4 |

[a] Engineered restriction sites underlined

[b] 5'end of *bs2* underlined

[c] 5'end P-*msp* underlined

[d] 6x-His fragment lower case

[e] 3'end P-*msp* underlined

[f] *msp* region DNA underlined

**Plasmid sequence alignments**

pTD1 isolated from *T. denticola* ATCC 33520 (Genbank M87856; 1992)

pTD1 isolated from *T. denticola* ATCC 33520 (Fenno lab stock; 2024)

pTD1 isolated from *T. denticola* clinical isolate CF219 (J Clin Microbiol. 25(11):2230-2).

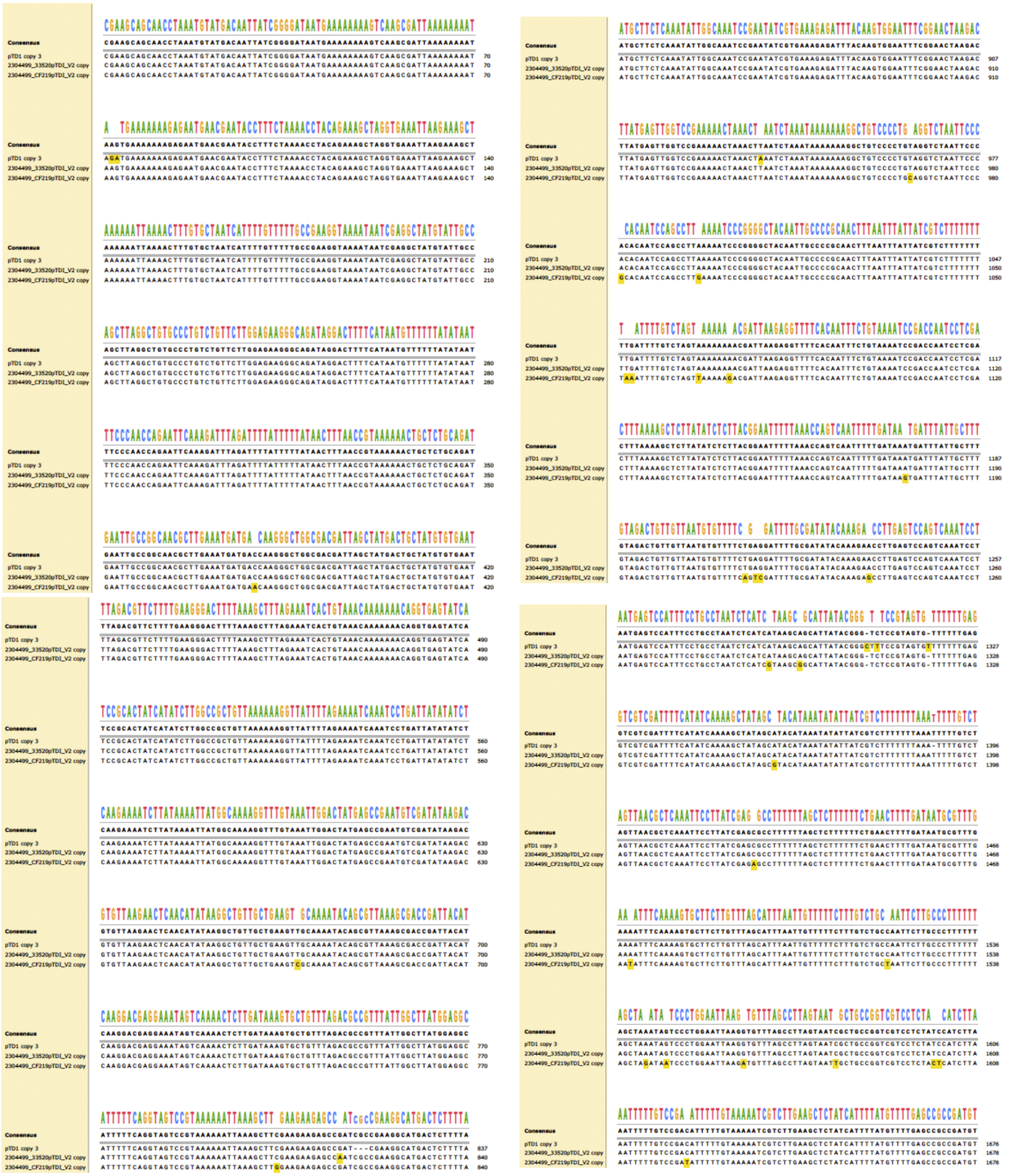

(continued on next page)

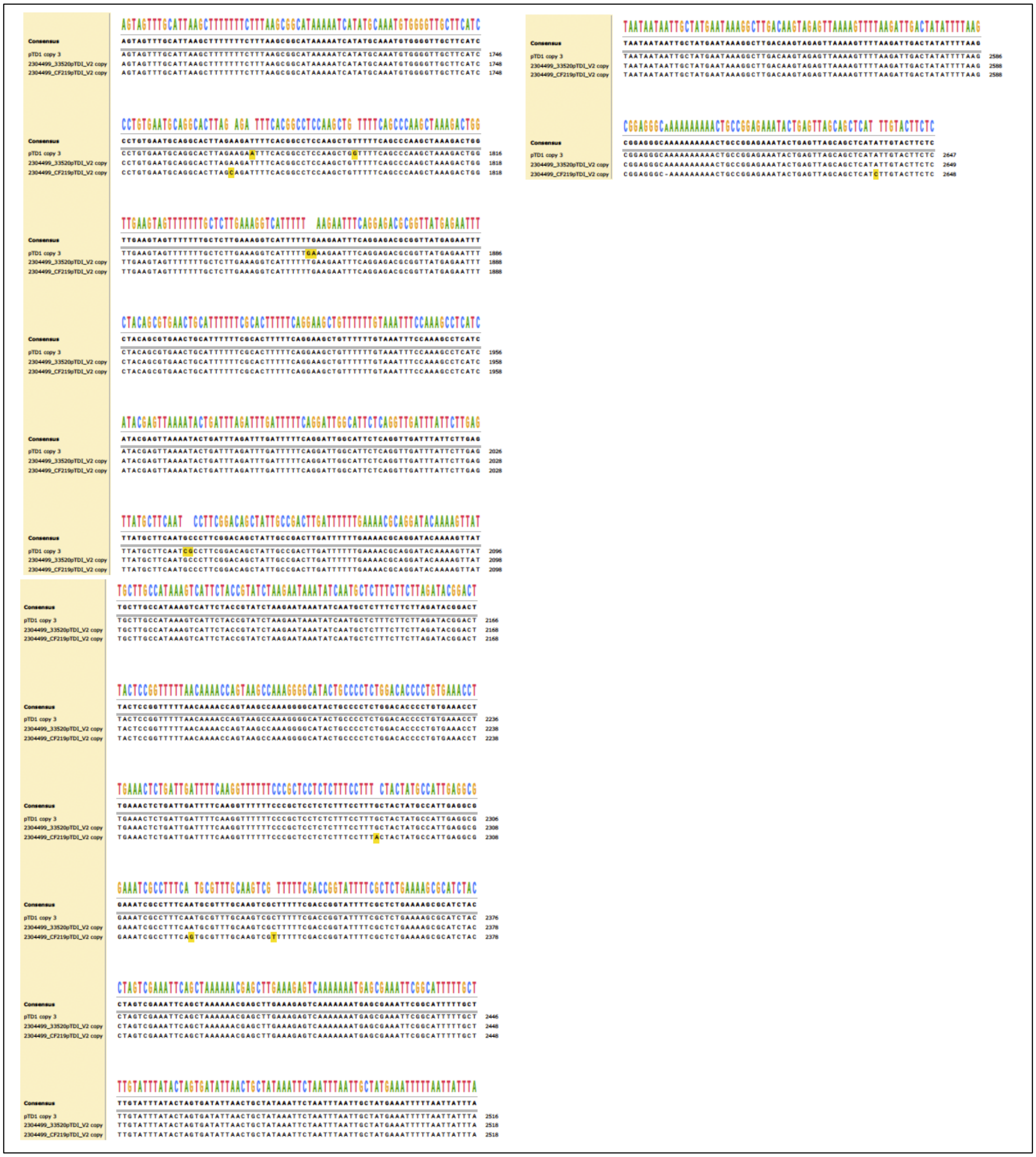
